# In-gel refolding allows fluorescence detection of fully denatured GFPs after SDS-PAGE

**DOI:** 10.1101/2024.10.02.615947

**Authors:** Misa Shiratori, Rio Tsuyuki, Miwako Asanuma, Saki Kawabata, Hiromasa Yoshioka, Kenji Ohgane

**Affiliations:** Department of Chemistry, Faculty of Science, Ochanomizu University, 2-1-1 Otsuka, Bunkyo-ku, Tokyo 112-8610, Japan; Chemical Biology Research Group, RIKEN Center for Sustainable Resource Science, Wako, Saitama 351-0198, Japan; Institute for Human Life Science, Ochanomizu University, 2-1-1 Otsuka, Bunkyo-ku, Tokyo 112-8610, Japan

**Keywords:** GFP, in-gel fluorescence, in-gel refolding, SDS-PAGE, cyclodextrin

## Abstract

Green fluorescent proteins (GFPs) have been widely used as fusion tags, especially to visualize subcellular localization and dynamics of the fused partner proteins. Also, GFPs serve as fluorescent tags in size-exclusion chromatography and native-PAGE, facilitating the evaluation of expression levels and quality of the expressed fusion proteins. However, the fluorescent detection of GFPs is generally incompatible with denaturing SDS-polyacrylamide gel electrophoresis (PAGE), where the samples are heat-denatured before loading. Accordingly, detecting GFP-fused proteins after SDS-PAGE usually relies on western blotting with anti-GFP antibodies. To enable in-gel fluorescence detection of SDS-PAGE-separated GFPs, some protocols employ mild denaturing conditions to keep the GFPs intact. However, such mild denaturation sometimes results in partial denaturation of the proteins and irregular electrophoretic mobility that is not proportional to their molecular weights. Here, we demonstrate that the fully denatured GFPs can be refolded within the gel by cyclodextrin-mediated removal of SDS in the presence of 20% methanol, enabling the in-gel fluorescence detection of the GFP-fused proteins. The protocol is compatible with subsequent total protein staining and western blotting. Although future studies are needed to clarify the scope and generality, the technique developed here would provide a simple, time- and cost-effective alternative to the immunodetection of GFPs.

**Highlights:**

- Fully denatured GFPs could be in-gel refolded to enable fluorescence detection.
- ? cyclodextrin effectively removed SDS to assist in the in-gel refolding of GFPs.
- GFPs amenable for in-gel refolding include GFPuv, EGFP, turboGFP, and sfGFP.
- Compatible with total protein staining and western blotting.
- The in-gel refolding procedure was successfully applied to two turboGFP-fusion proteins.

**Graphical abstract:** 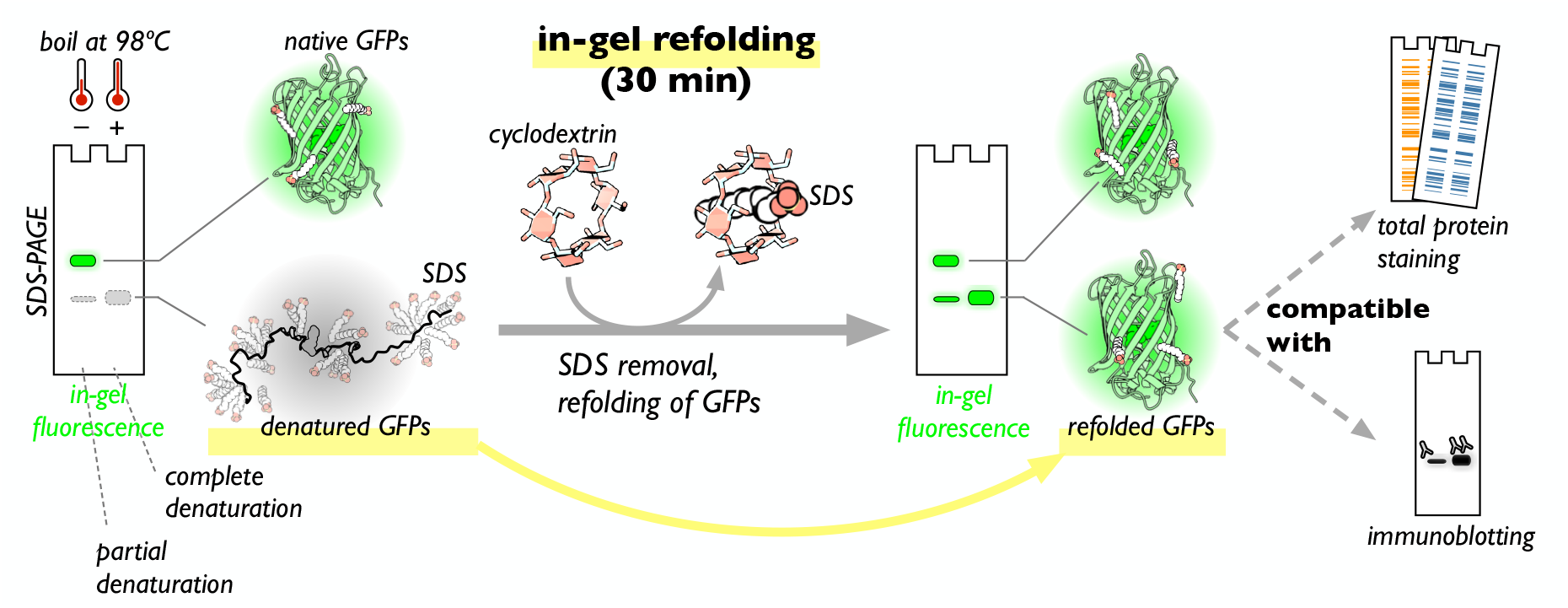

## 1. Introduction

Green fluorescent proteins (GFPs) and other related fluorescent proteins (FPs) are now essential fusion-protein tags that enable visualization of subcellular localization and dynamics of the fused partner proteins in live cells. Also, GFPs serve as rapid and sensitive fluorescent tags for chromatography and electrophoresis, including fluorescent-detection size-exclusion chromatography (FSEC) and high-resolution clear-native PAGE [1-3]. Thus, GFP tagging made it easy to screen expression level, monodispersity, thermostability, and correct subcellular localization of the fusion proteins, greatly facilitating structural studies of proteins, especially membrane proteins [4-8].

GFP fluorescence is normally lost after standard SDS-PAGE. In SDS-PAGE, the protein sample is denatured before loading onto the gel, by boiling in the Laemmli sample buffer which contains SDS and a reducing agent like DTT, making it unsuitable to detect GFP fluorescence after SDS-PAGE. However, by exploiting the high stability of the folded GFPs, in-gel fluorescence detection is possible if only the protein sample is not boiled or incubated at a lower temperature in the sample buffer, to avoid denaturation of GFPs while denaturing most of the other proteins [9,10]. Such protocols enable time- and cost-effective detection of GFP-tagged proteins even at endogenous expression levels and can offer better quantification with a broader dynamic range [10]. It potentially overcomes problems often observed for immunoblotting, including heterogeneous protein transfer, a limited dynamic range of enhanced chemiluminescence detection, and substantial cost due to the antibodies and other reagents. One of the drawbacks of this protocol is that the partial denaturation gives anomalous electrophoretic mobility that deviates from expected mobility based on molecular weight, due to the folded structure of GFPs. Another is that multiple bands can be observed for folded and unfolded GFP-fused proteins even under milder denaturation condition, requiring appropriate controls and attention while interpreting the results [10].

In principle, removing denaturants can refold proteins [11]. Some enzymes consisting of one polypeptide chain were shown to refold after denaturing SDS-PAGE by simply washing the gels with appropriate buffers for 1-20 hours, where protein-bound SDS spontaneously diffuses out of the gels [12, 13]. However, the diffusion of the electrophoresed proteins during the long wash makes the detection less sensitive and results in diffuse bands. Later studies identified that 25% isopropanol is effective in removing SDS and refolding nucleases in the gels [14], and a similar condition is now adopted for in-gel detection of a small and stable luciferase, NanoLuc (In-Gel Detection System of NanoLuc, Promega). We reasoned that an appropriate condition for refolding GFPs would enable in-gel detection of GFP-fused proteins after denaturing SDS-PAGE even with boiled, fully denatured samples. There exist several precedents that successfully refolded GFPs after SDS-PAGE. For refolding of in-vitro transcribed GFP, a mildly boiled sample (80ºC for 5 min) was resolved on SDS-PAGE, and the gel was shaken in the standard transfer buffer (25 mM Tris, 192 mM Glycine, 20% MeOH) for 30 min to successfully detect GFP fluorescence [15]. In another example, a fusion protein with mTurquoise2, a rapidly maturing monomeric cyan fluorescent protein (FPbase ID: B4SOW), was partially refolded after denaturing SDS-PAGE by washing the gel in distilled water for 30 min [16, 17]. However, optimization of the protocol has not been reported and the generality of the technique remains unclear.

In the present study, we report an optimized procedure for in-gel refolding of GFPs after SDS-PAGE of boiled, denatured GFP-tagged proteins. Even with the fully denatured samples, an additional short refolding step can resume the fluorescence signal, allowing in-gel fluorescence detection of the GFPs. The protocol is compatible with subsequent total protein staining and western blotting with antibodies. This protocol would offer a time- and cost-effective option for the current common western blotting workflows and provide a complementary alternative to the “non-boiling” SDS-PAGE protocol for in-gel GFP detection [9, 10].

## 2. Material and methods

### 2.1 Materials

The pGLO plasmid, encoding GFPuv under the control of AraBAD promoter, was obtained from Bio-RAD. pEGFP-N1, encoding EGFP under the control of CMV promoter was from Clontech. pReceiver-EGFP (M02Rx) was from GeneCopoeia. Both α- and β-cyclodextrin were purchased from FUJIFILM Wako.

### 2.2 Site-directed mutagenesis and MEGAWHOP cloning

We constructed pGLO-high-sfGFP by introducing a high-copy number mutation (svir002 mutation) into the replication origin [18] and substituting GFPuv with superfolder GFP (sfGFP) as described below [19]. The site-directed mutagenesis was performed using a pair of partially overlapping primers and Phusion DNA polymerase (New England Biolabs) [20]. Briefly, following PCR amplification with pGLO plasmid and a pair of primers (fwd: *CCAAATACTGT<u>T</u>CTTCTAGTG*TAGCCGTAG, rev: *CACTAGAAG<u>A</u>ACAGTATTTGG*TATCTGCG, underlines indicate the C to T mutation and the italicized bases denote the overlapping region), DpnI digestion removed the template plasmid, and the product was directly used to transform competent Quick DH5α cells (TOYOBO) to obtain pGLO-high plasmid after miniprep (Bio-RAD). The GFPuv sequence in the pGLO-high plasmid was replaced with a megaprimer-based whole plasmid amplification (MEGAWHOP) cloning [21, 22]. Briefly, PCR amplification was performed with the pGLO-high plasmid and a complementary double-stranded DNA containing sfGFP as megaprimers (synthesized by Thermo Fisher Scientific GeneArt, for sequence see Supplementary Methods) using KOD-plus-DNA polymerase. Following DpnI digestion to remove the template plasmid, the reaction product was used to transform competent Quick DH5α cells (TOYOBO), yielding pGLO-high-sfGFP plasmid. A monomerizing mutation, V206K, was introduced to pGLO-high-sfGFP similarly as described above using Phusion DNA polymerase (New England Biolabs) and the following pair of primers (fwd: *CCTGTCGACACAATCTAAGC*TTTCGA<u>AAG</u>ATC, rev: *G<u>CTT</u>AGATTGTGTCGACAGG*TAATGGTTGTCTG, the underlines indicate the V206K mutation and the italicized bases denote the overlapping region required for MEGAWHOP). The integrity of the plasmids was confirmed by sequencing (Sanger sequencing at FASMAC and PlasmidEZ at Azenta).

A plasmid vector for FLAG-EGFP-OSBP expression was also constructed in two steps. First, the MEGAWHOP cloning with a synthesized megaprimer containing the human OSBP coding sequence (synthesized by Thermo Fisher Scientific GeneArt, for sequence see Supplementary Methods) inserted the OSBP sequence into pReceiver-EGFP vector (GeneCopoeia). The subsequent site-directed mutagenesis with Phusion DNA polymerase inserted the FLAG tag (DYKDDDDK) and a Gly after the first methionine in the EGFP. The following set of primers was used (fwd: gactacaaggatgacgatgacaagggcGTGAGCAAGGGCGAGGAGCTGTTC and rev: gcccttgtcatcgtcatccttgtagtcCATGGTACCGAATTCTTCGAAGATC, the lower cases denote the overlapping region, which corresponds to DYKDDDDKG sequence.

### 2.3 Expression of GFPs in DH5α and protein harvest

For bacterial expression of sfGFP, we transformed DH5α competent cells with pGLO-high-sfGFP, and expression was induced by adding 0.1% L-arabinose during overnight culture at 37ºC in LB medium supplemented with ampicillin. To express GFPuv or sfGFP-V206K, pGLO or pGLO-high-sfGFP-V206K was used instead, respectively. The overnight culture was centrifuged at 14000×g for 5 min, and the pellet was resuspended in Tris-buffered saline (TBS) and disrupted by sonication on ice. After centrifugation at 14000×g for 5 min to remove cell debris, the supernatant was collected and processed for SDS-PAGE.

### 2.4 Cell lines and cell culture

HEK293 cell line (ATCC) was cultured in DMEM (high glucose) supplemented with 10% (v/v) FBS, penicillin/streptomycin solution (100 units/ml and 100 μg/ml, respectively, from Nacalai tesque). A clonal HEK293 cell line expressing FLAG-EGFP-OSBP was established by selection with G418 and the limited dilution procedure. The clones of stable HEK293 cell lines expressing NPC1-NTDtail-FLAG-tGFP or SM-FLAG-tGFP under the control of CMV promoter were previously described [23-25]. These stable cell lines were maintained in DMEM (high glucose) supplemented with 5% (v/v) FBS, penicillin (100 units/ml), and streptomycin (100 μg/ml).

### 2.5 SDS-PAGE

The lysate was mixed with 0.25 vol of 5×Laemmli sample buffer (10% SDS, 312.5 mM Tris-HCl pH 6.8, 0.025% bromophenol blue, 500 mM DTT, and 40% (v/v) glycerol) and heated at the indicated temperatures for 5 min. The samples were frozen at -20ºC until use. SDS-PAGE was carried out on an Easy Separator electrophoresis system (FUJIFILM-Wako) with myPowerII 500 power supply (AE-8155, ATTO) in gels (90 mm × 85 mm × 1mm thickness) comprised of 12% or 7.5% resolving gel and 4% stacking gel. SDS-PAGE was performed in standard SDS-PAGE running buffer (25 mM Tris, 192 mM glycine, 0.1% SDS) with a constant current of 20 mA per gel for 80 min, or 40 mA for 40 min, at room temperature. The gel after SDS-PAGE was briefly rinsed with deionized water and a fluorescence image was acquired on an iBright FL 1500 imaging system (Thermo Fisher Scientific).

### 2.6 In-gel refolding of GFPs

Following SDS-PAGE, the gels were shaken in the optimized renaturation buffer (1% αCD, 25 mM Tris, 192 mM Gly, and 20% (v/v) MeOH) for 30 min at room temperature. The gels were briefly rinsed with deionized water and fluorescence images were acquired on an iBright FL 1500 imaging system (Thermo Fisher Scientific). The following setting was used; Ex 455-485 nm and Em 508-557 nm for GFPs, resolution 3×3, exposure time 5 msec to 30 sec depending on the samples. Fluorescence signal from the gels was quantified with ImageJ and an ImageJ macro, band peak quantification tool [26]. The apparent molecular weight was estimated from the gel images by using BLUE Star PLUS prestained protein ladder (Nippon Genetics) and an ImageJ macro [27].

### 2.7 Total protein staining

Total proteins in the gels were stained with SYPRO Orange gel stain (Thermo Fisher Scientific). Briefly, the gel was shaken in 7.5% (v/v) acetic acid (AcOH) in water supplemented with 1/5000 SYPRO Orange stock solution at room temperature for 30 min. The stained gels were briefly rinsed with 7.5% AcOH and imaged on an iBright FL1500 imaging system (SYPRO Orange: Ex 490-520 nm / Em 568-617 nm).

### 2.8 Western blotting

Proteins were transferred from the gel after in-gel renaturation, which was equilibrated with a standard Towbin transfer buffer (25 mM Tris, 192 mM Gly, 20% (v/v) MeOH) for 10 min, to a PVDF membrane in a Mini-protean tetra wet transfer system (Bio-Rad) with myPowerII 500 power supply (AE-8155, ATTO). The transfer was performed with a constant voltage of 20 V (max 90 mA) for 16 h at 4ºC or 40 V (max 250 mA) for 2h. The membranes were blocked in 1% BSA in TBS supplemented with 0.1% Tween 20 (TBST) for 30 min at room temperature or overnight at 4ºC. The membranes were incubated with mouse monoclonal anti-FLAG M2 (Sigma-Aldrich, F1804) or rabbit polyclonal anti-GFP (life technologies, A11122, 1/5000) antibodies diluted in 10% Can Get Signal solution 1 (TOYOBO) for 1-2h. After extensive washing with TBST, the membranes were incubated with a goat polyclonal HRP-conjugated anti-mouse IgG (Millipore, 12-349, 1/5000) or goat polyclonal HRP-conjugated anti-rabbit IgG (Cell Signaling, 7074S, 1/5000) antibodies diluted in 10% Can Get Signal solution 2 (TOYOBO) for 1 h. After washing with TBST, the blot was visualized by enhanced chemiluminescence [28] and imaged on an iBright FL 1500 imaging system.

### 2.9 Protein precipitation

#### 2.9.1 Chloroform-methanol precipitation

The lysate (50 µL) on ice was added MeOH (150 µL), chloroform (37.5 µL), and water (100 µL) followed by vortexing at each addition. The mixture was centrifuged at 14,000×g for 5 min and the upper aqueous phase was removed. Additional MeOH (150 µL) was added to the tube, vortexed, and centrifuged at the same condition. The supernatant was removed, and the pellet was air-dried for 5 minutes. The pellet was re-solubilized with 1×Laemmli sample buffer (50 µL) for 15 minutes at room temperature and 5 minutes at the indicated temperature.

#### 2.9.2 Acetone precipitation

The lysate (50 µL) on ice was added cold acetone (200 µL) and incubated at -20ºC for 1 hour. The tube was centrifuged at 14,000×g for 10 min and the supernatant was removed. The pellet was air-dried for 20 minutes and re-solubilized with the 1×Laemmli sample buffer (50 µL) for 15 minutes at room temperature and 5 minutes at the indicated temperature.

#### 2.9.3 Trichloroacetic acid (TCA) precipitation

The lysate (50 µL) on ice was added TCA (99% (w/v) aqueous solution, 12.5 µL) and incubated at 0ºC for 5 minutes. The tube was centrifuged at 14,000×g for 10 min and the supernatant was removed. Cold acetone (125 µL) was added to the tube and centrifuged at the same condition, and this wash step was repeated. The pellet was air-dried for 20 minutes and re-solubilized with the 1×Laemmli sample buffer (50 µL) for 15 minutes at room temperature and 5 minutes at the indicated temperature.

## 3. Results and discussion

### 3.1 Optimization of in-gel refolding protocol for GFPs

To refold denatured GFPs in SDS-PAGE gel, SDS needs to be removed from the gel. We hypothesized that cyclodextrins and organic solvents could effectively remove SDS. Cyclodextrins (CDs) have been used to remove SDS from proteins via cyclodextrin-SDS inclusion complex formation [29, 30], and isopropanol was applied to refold some enzymes after SDS-PAGE [14]. For refolding of GFPs, we selected Tris-glycine (Tris-Gly) buffer (approximately pH 8.3) as a buffer, considering that GFPs are more stable at slightly basic conditions and their fluorescence is quenched at acidic conditions [10, 31]. An additional advantage would be that this buffer can be easily prepared from the standard 10×transfer buffer for western blotting.

Using lysate of *E. coli* expressing sfGFP (FPbase ID: B4SOW) as a model [19], we tested the ability of Tris-Gly buffers with various organic solvents to refold fully denatured sfGFP in SDS-PAGE gel (**Fig. 1A**). Lysates containing sfGFP were mixed with Laemmli sample buffer, kept at 98ºC or 0ºC for 5 min, and resolved by standard SDS-PAGE. The gels were then shaken in the indicated buffers and the in-gel fluorescence signal was detected. As we will discuss later, the denatured sfGFP migrated at an apparent molecular weight of 26 kDa and the non-denatured sfGFP at 40 kDa. Among the organic solvents tested, isopropanol (iPrOH) gave the best refolding efficiency (69% fluorescence relative to the non-boiled sample), followed by acetonitrile (MeCN, 17% fluorescence). Methanol (MeOH), commonly included at 10-20% in the transfer buffer for Western blotting, or ethanol (EtOH) did not assist in SDS removal over spontaneous diffusion in the Tris-Gly buffer nor support the refolding of sfGFP. Next, we varied the concentration of isopropanol from 0% to 25% and found that concentrations of more than 20% enhanced refolding efficiency at 20 min (**Fig. 1B**).

**Figure 1.**
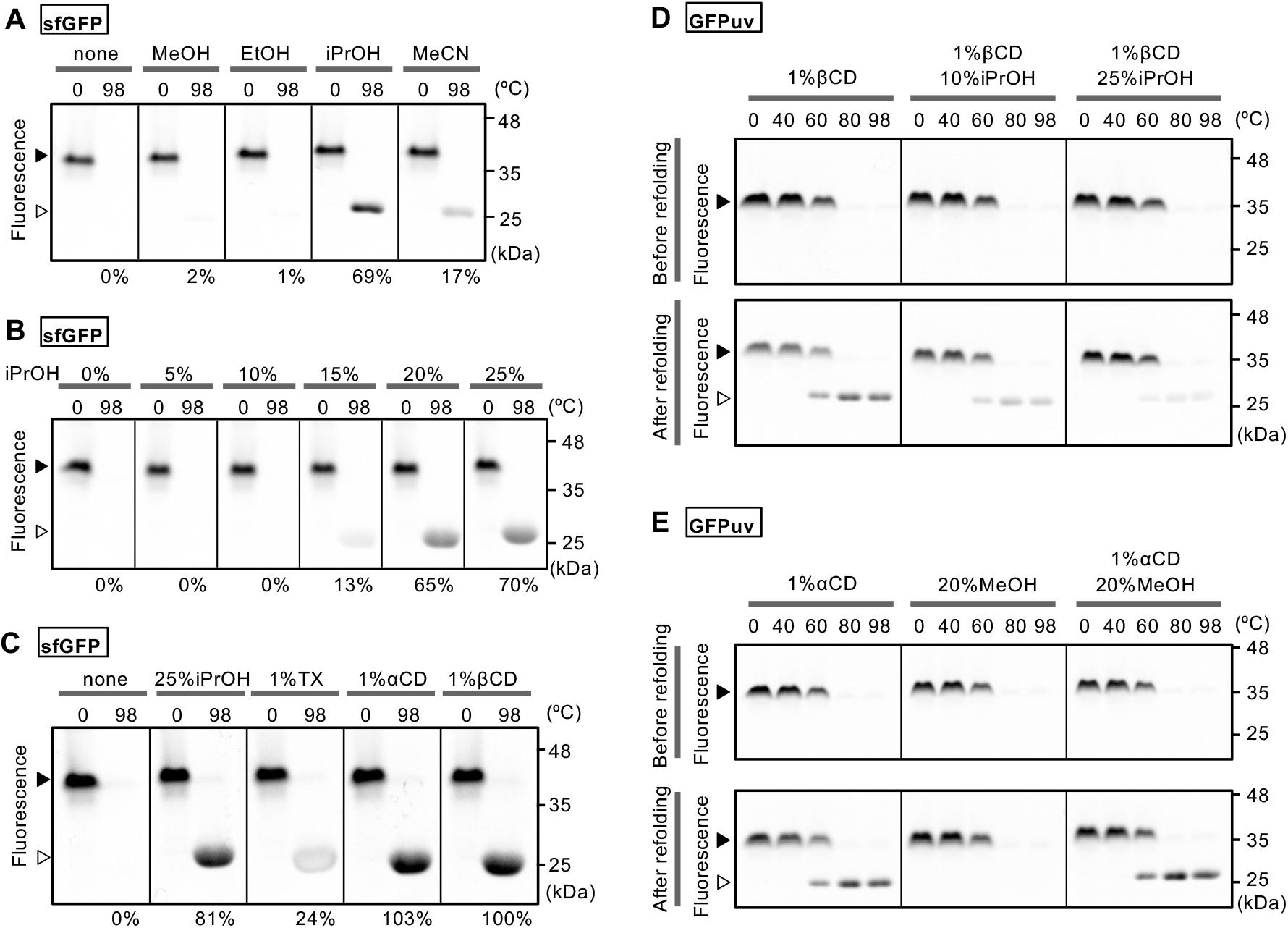
Optimization of the in-gel refolding condition for heat-denatured sfGFP. **A**. Effect of organic solvents on the in-gel refolding of heat-denatured sfGFP. Different organic solvents were tested for their efficiency in assisting the in-gel refolding. After SDS-PAGE, the gel was cut into strips and each strip was shaken in the Tris-Gly buffer with 20% (v/v) of the indicated organic solvents for 20 min at room temperature. GFP fluorescence from the set of strips was imaged simultaneously and shown with the same contrast. The values below the gel images denote fluorescence intensity from the heat-denatured band (98ºC) relative to the non-denatured band (0ºC) within the same strip of the gel. The filled and open triangles represent the non-denatured 40 kDa band and the denatured 26 kDa band, respectively. **B**. Optimal concentration of iPrOH for in-gel refolding of heat-denatured sfGFP. The experiment was performed as described in **A**. The filled and open triangles represent the non-denatured 40 kDa band and the denatured 26 kDa band, respectively. **C**. Effect of Triton X-100 (TX) and CDs on the in-gel refolding efficiency of heat-denatured GFPuv, performed as in **A. D**. Effect of iPrOH in the presence of 1% βCD on the refolding efficiency and fluorescence intensity. The lysate was heated for 5 min at the indicated temperature in the Laemmli sample buffer and resolved on SDS-PAGE. The gel after SDS-PAGE was first imaged for GFP fluorescence and then cut into strips. The strips were shaken in the Tris-Gly buffer with the indicated additives for 20 min, and GFP fluorescence was imaged simultaneously and shown with the same contrast. The filled and open triangles represent the non-denatured 35 kDa band and the denatured 26 kDa band, respectively. **E**. In-gel refolding of GFPuv under the optimized refolding condition (Tris-Gly buffer with 1% αCD and 20% MeOH) or the buffers containing either 1% αCD or 20% MeOH. The filled and open arrowheads represent the non-denatured 35 kDa and the denatured 26 kDa band, respectively.

Next, we tested if cyclodextrins could remove SDS to assist in in-gel refolding of fully denatured sfGFP. When the gel was washed with Tris-Gly buffer containing 1% α-cyclodextrin (αCD) or 1% β-cyclodextrin (βCD), both cyclodextrins enhanced refolding of sfGFP, yielding nearly quantitative recovery of the fluorescence (**Fig. 1C**). Although both αCD and βCD performed similarly, we selected αCD as the optimal additive for in-gel refolding of GFP, considering the higher solubility of αCD (soluble at >10% in water) over βCD (1% in water). We also tested less-denaturing detergents including Triton X-100 (**Fig. 1C**), sodium deoxycholate (data not shown), and CHAPS (data not shown), which might competitively replace SDS on the proteins. However, they showed no improvement in assisting in refolding sfGFP. This is consistent with the inability of Triton X-100 to induce refolding of DNase I [13].

As sfGFP is known for its high thermal stability and fast folding kinetics, further optimization was performed with a less stable variant GFPuv (FPbase ID: B28N7), also known as cycle 3 variant or αGFP, expressed in *E. coli*. 1% βCD worked fine for GFPuv as well, regenerating fluorescence from the heat-denatured samples (**Fig. 1D**). While both iPrOH and cyclodextrins performed well for sfGFP, the addition of iPrOH had a negative impact on the refolding of GFPuv, probably reflecting the less stable nature of GFPuv compared to sfGFP. Additionally, we noted that the refolding buffer without alcohols led to reduced GFPuv fluorescence from the non-denatured band at 35 kDa (**Fig. 1D**). Probably this diminished fluorescence was due to diffusion of the proteins from the gels. The presence of alcohols played a role in preventing the diffusion of proteins from the gels during the refolding. In contrast to the iPrOH, 20% MeOH did not interfere with the refolding of GFPuv (**Fig. 1E**). Overall, we propose 1% αCD in Tris-Gly buffer containing 20%MeOH as an optimized buffer for in-gel refolding of GFPs.

We next tested if EGFP, a widely used GFP (FPbase ID: R9NL8), expressed in mammalian cell culture is also amenable for in-gel refolding. The fully denatured EGFP in cell lysate from transiently transfected HEK293 cells was separated on the standard SDS-PAGE and subjected to in-gel refolding before fluorescence detection. Consistent with the data shown above, shaking in the Towbin transfer buffer (Tris-Gly, 20% MeOH) for 30 min was ineffective in assisting in-gel refolding of EGFP and the addition of 1% αCD almost completely regenerated its fluorescence (**Fig. 2**). The use of 25% isopropanol instead of 20% MeOH was less effective and showed slightly lower refolding efficiency, again reflecting the less efficient folding of EGFP than sfGFP. After in-gel refolding, the gel was stained for total proteins with SYPRO Orange stain, supporting the compatibility with total protein staining.

**Figure 2.**
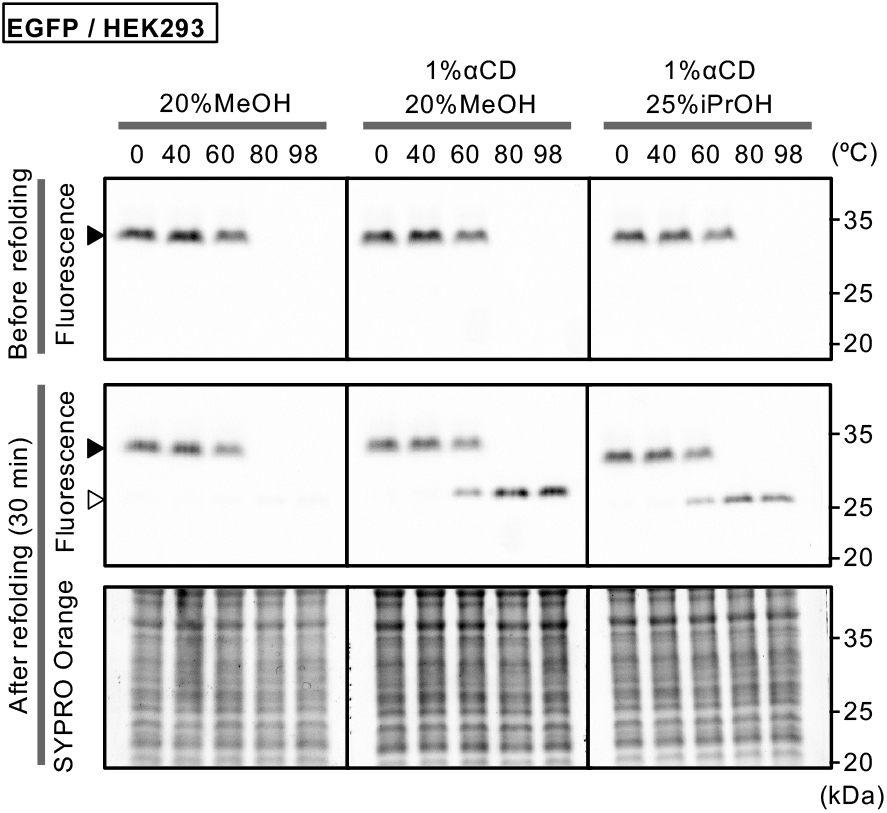
In-gel refolding of EGFP expressed in HEK293 cells. Aliquots of cell lysate from HEK293 cells transiently transfected with pEGFP-N1 were mixed with Laemmli sample buffer and heated at the indicated temperature for 5 min before SDS-PAGE. The gel was imaged before and after the in-gel refolding at the indicated conditions (30 min). The gel was also stained with SYPRO Orange dye for total protein. The filled and open triangles represent the non-denatured 33 kDa band and the denatured 27 kDa band, respectively.

For all of sfGFP, GFPuv, and EGFP, non-denatured GFP exhibited slower migration (apparent molecular weight of 40 kDa) than the fully denatured, boiled GFP (apparent molecular weight of 27 kDa, consistent with their amino acid sequences), as previously observed [10]. A likely explanation would be that the folded structure of the non-denatured species binds less SDS and therefore migrated slowly in the gel. However, it is also possible that the GFPs tested above tend to form a dimer, and the slower migrating species represent dimers. So, we tested this possibility by using a monomeric sfGFP variant harboring V206K mutation [32]. The monomeric V206K variant also exhibited a similar pattern (39 kDa non-denatured band and 24 kDa denatured band) (**Fig. 3**), arguing against the observed slower migrating species representing dimeric species. Thus, the non-boiling SDS-PAGE protocol, which requires intact GFP, can result in irregular electrophoretic mobility due to the folded structure of GFP [10].

**Figure 3.**
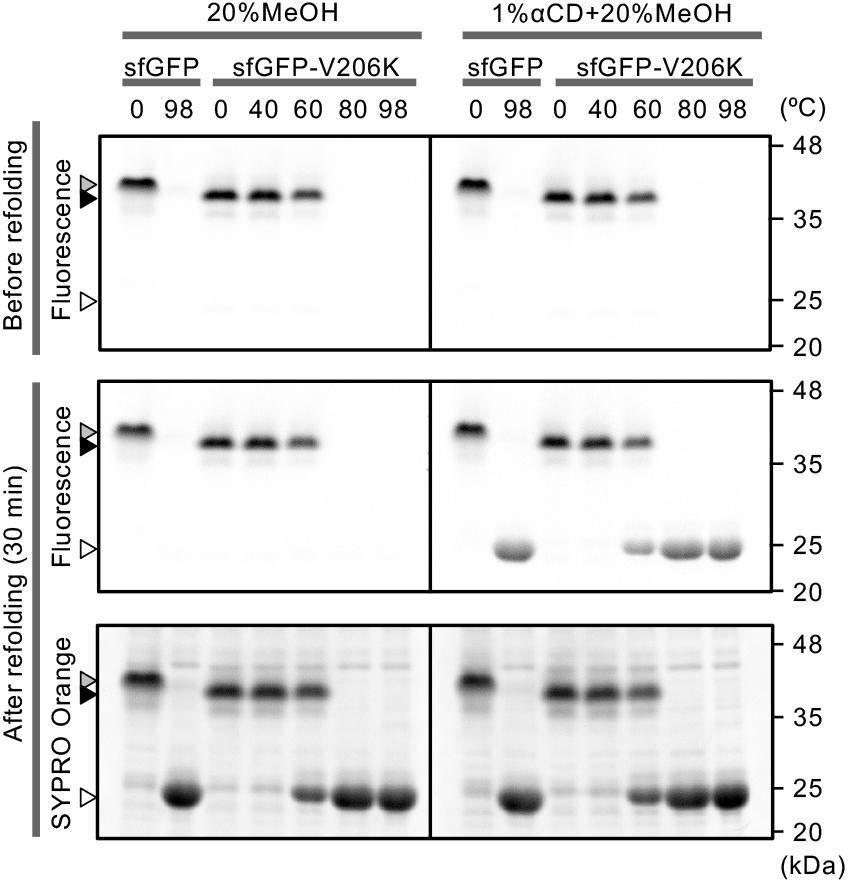
In-gel refolding of sfGFP and sfGFP-V206K, a monomerizing mutant. Lysates of E. coli expressing each sfGFP variant were heated at the indicated temperature for 5 min in the Laemmli sample buffer and resolved on the SDS-PAGE. The gel was renatured in the Tris-Gly buffer with the indicated additives for 30 min with gentle shaking and GFP fluorescence was imaged. After renaturation, the gel was stained in the SYPRO Orange solution containing 7.5% acetic acid for 30 min before fluorescence imaging. The gray, black, and white arrowheads represent 41 kDa, 39 kDa, and 24 kDa bands, respectively.

Precipitation of proteins with trichloroacetic acid (TCA), acetone, and chloroform-methanol is a useful technique especially to concentrate dilute protein solution or to remove undesired low molecular weight contaminants. TCA precipitation has been shown to denature GFP and is incompatible with the in-gel fluorescent detection after the non-boiling SDS-PAGE [10]. Next, we tested whether GFP denatured during the common protein precipitation methods could be refolded after SDS-PAGE and enable in-gel fluorescence detection. The lysate from *E. coli* expressing sfGFP was subjected to TCA precipitation, acetone precipitation, or chloroform-methanol precipitation, and the precipitated proteins were re-solubilized with Laemmli sample buffer at the indicated temperatures (**Fig. 4**). Without refolding, TCA- or chloroform-methanol-precipitated samples showed no fluorescence, replicating the previous observation [10]. After the refolding procedure, in-gel fluorescence was observed for all the precipitated samples, including TCA precipitated samples re-solubilized at 98ºC. We also noted that the acetone precipitation did not completely denature sfGFP; the re-solubilized pellet at room temperature gave a fluorescent band without refolding probably due to the high stability of sfGFP and relatively mild denaturing effect of the acetone precipitation. Thus, our in-gel refolding procedure was also effective for sfGFP denatured during the precipitation/re-solubilization sequence.

**Figure 4.**
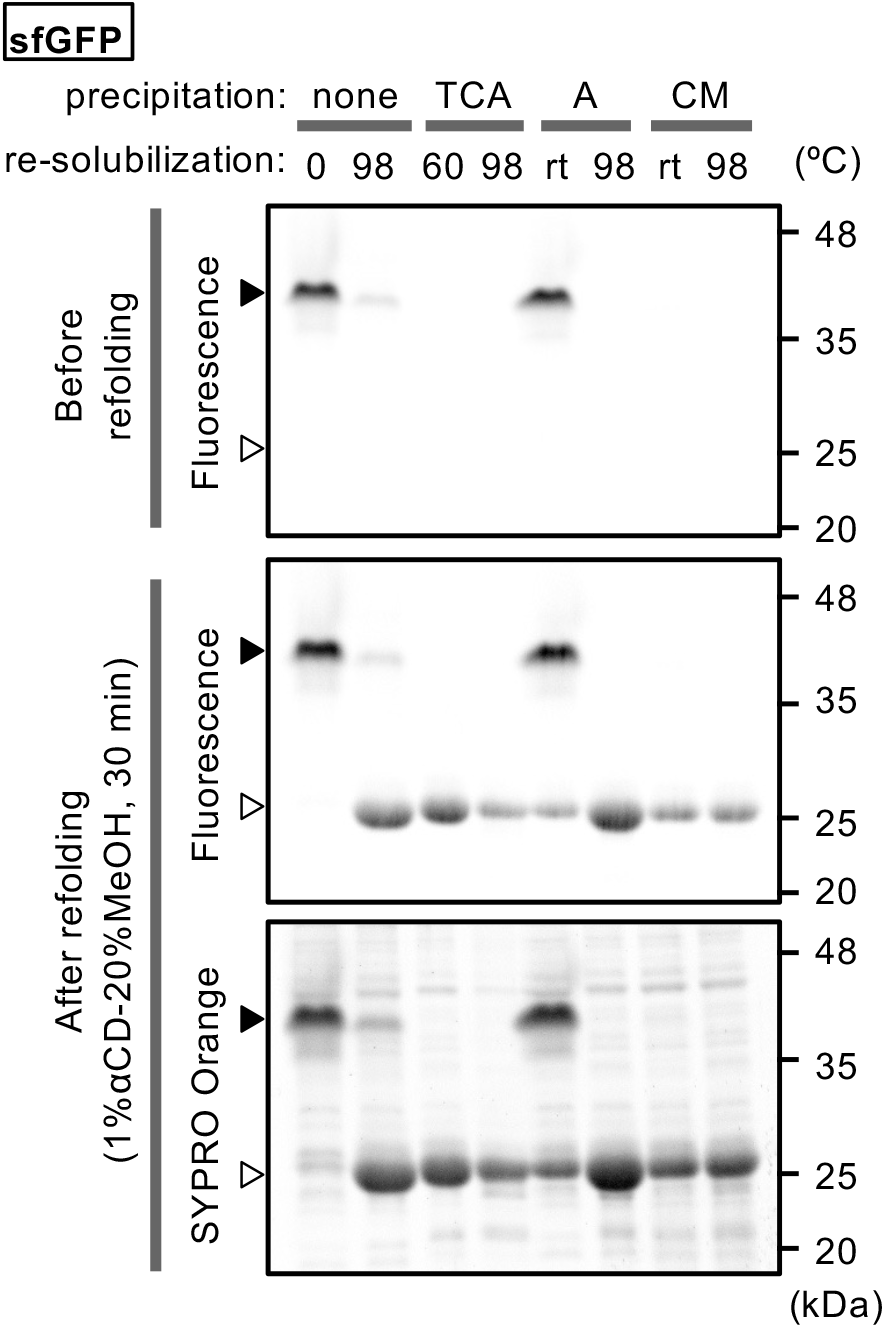
In-gel refolding and fluorescence detection of TCA-, acetone-, chloroform-methanol-precipitated sfGFP. The lysate (50 µL) containing sfGFP was precipitated with TCA, acetone (A), or chloroform-methanol (CM), and the precipitate was resolubilized in 50 µL of 1×Laemmli sample buffer at room temperature for 15 min then at the indicated temperature for 5 min. The samples were resolved on standard SDS-PAGE, and in-gel fluorescence and refolding were performed as described above. The filled and open arrowheads denote the non-denatured 42 kDa band and the denatured 26 kDa band, respectively.

### 3.2 Application to GFP-fused proteins expressed in mammalian cells

Next, we focused on GFP fusion proteins expressed in human cell lines to further clarify the utility of the in-gel refolding protocol. We tested a fragment of Niemann-Pick type C1 (NPC1) membrane protein fused with turboGFP (tGFP, FPbase ID: A315M), a rapidly maturing GFP from a marine copepod. The full-length NPC1 protein, a lysosomal 13-pass membrane protein with 1278 amino acid residues, is known to disappear after boiling due to irreversible aggregation, as is often observed for multi-pass membrane proteins [4]. Therefore, we used a fragment of NPC1 (Δ306-1256) which contains a heavily glycosylated N-terminal domain (NTD) facing the lysosomal lumen, the first transmembrane domain, and its C-terminal tail with lysosomal targeting motif (NPC1-NTDtail) [23, 24]. The GFP used here is turboGFP, characterized by its fast maturation in living cells despite being thermally less stable than EGFP or sfGFP [33-35]. In-gel fluorescence of NPC1-NTDtail-tGFP revealed the doublet band at 70 and 73 kDa for the samples heated at 40ºC or lower temperatures, but the corresponding bands were undetectable for the other samples heated at more than 80ºC due to heat-denaturation of the GFP (**Fig. 5A, top**). The observation of the doublet band was consistent with our previous observation; this protein is a heterogeneously glycosylated membrane protein and gives a doublet band on immunoblots [23]. When the gel was shaken in the optimized refolding buffer for 30 min, an additional pair of doublet bands at 76 and 84 kDa became visible for the heat-denatured samples as well. This larger doublet band became visible even in the sample prepared without heating (0ºC) (**Fig. 5A, middle**), indicating that this fusion protein is likely relatively labile; more than half of the protein was denatured without heating.

**Figure 5.**
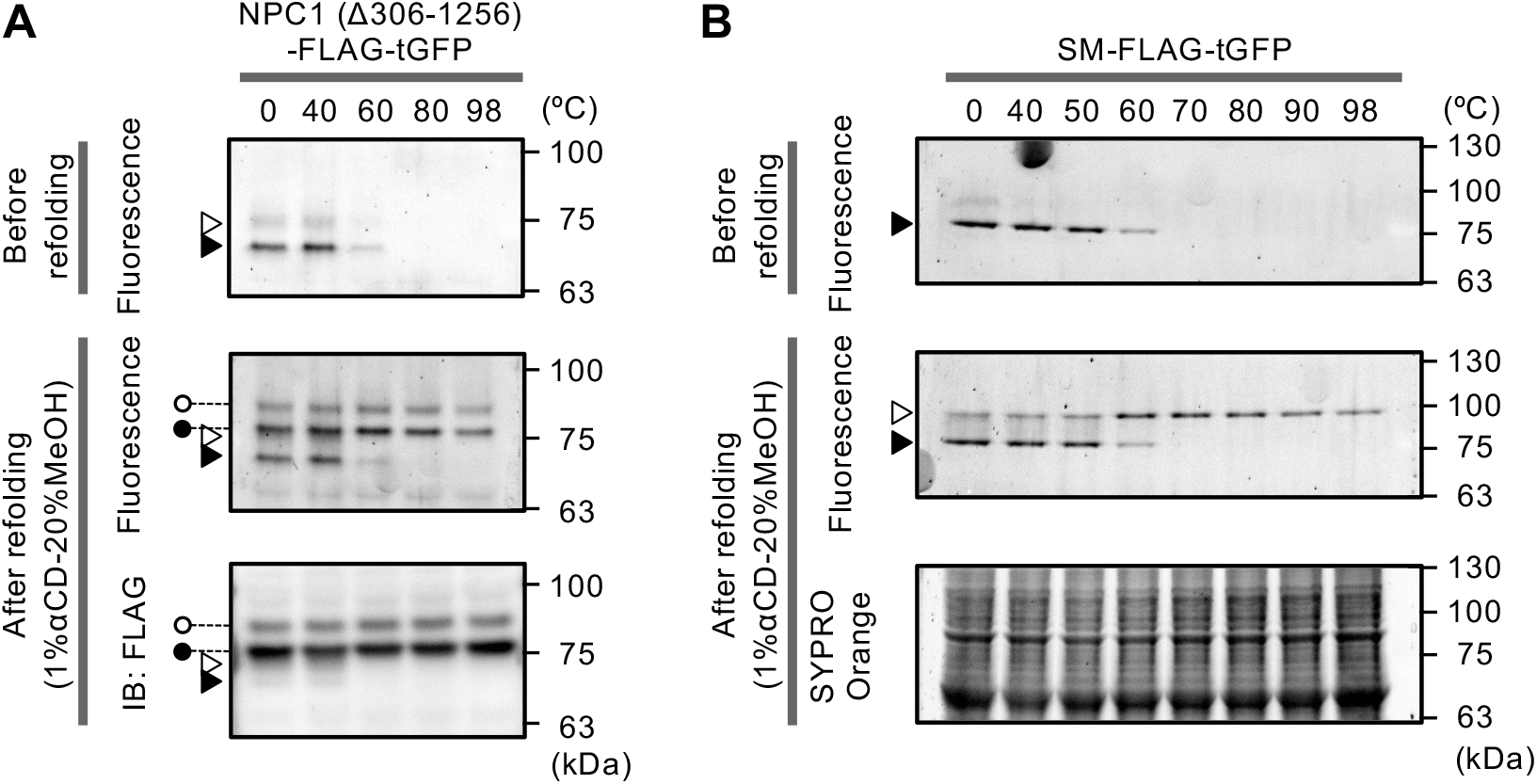
Application of the refolding procedure to tGFP-fusion proteins. **A**. In-gel refolding of a single-pass membrane protein, NPC1 (Δ306-1256)-FLAG-tGFP stably expressed in HEK293 cells. Aliquots of cell lysate mixed with Laemmli sample buffer were heated at the indicated temperature for 5 min and resolved on an SDS-PAGE. The GFP fluorescence was imaged before and after the in-gel refolding with the Tris-Gly buffer supplemented with 1% αCD and 20% MeOH (30 min). Proteins on the renatured gel were then transferred to a PVDF membrane, and the western blotting with anti-FLAG antibody was performed. The open and black arrowheads represent the non-denatured 73 and 70 kDa bands, respectively. The open and filled circles represent the denatured 84 and 76 kDa bands, respectively. **B**. In-gel refolding of SM-FLAG-tGFP stably expressed in HEK293 cells. The experiment was performed as described in A, except that the gel after refolding was stained with SYPRO Orange for total protein staining. The filled and open arrowheads represent the non-denatured 77 kDa band and the denatured 88 kDa band, respectively.

The renatured gel was also compatible with the standard Western blotting (**Fig. 5A, bottom**). Although SDS on the protein is partially removed during the refolding, SDS sufficient for electroblotting probably remains on the proteins. Additionally, we noted that fully denatured proteins (76 and 84 kDa) result in higher signal intensity on the immunoblot than the non-denatured bands at 70 and 73 kDa. The fully denatured species (76 and 84 kDa) were possibly more efficiently transferred to the PVDF membrane than the partially denatured species (70 and 73 kDa). Such behavior will need attention if the “non-boiling” SDS-PAGE is multiplexed with western blotting analysis, as the in-gel fluorescence and the western blotting data visualize different conformational states of the same fusion protein and give bands at different apparent molecular weights.

To further examine the scope of the refolding procedure, we tested if a membrane-associated enzyme squalene monooxygenase (SM) fused with tGFP could also be in-gel refolded [25, 36, 37]. Without boiling, a 77 kDa band was observed under fluorescence detection, which is smaller than the theoretical molecular weight of the fusion protein, 92 kDa (**Fig. 5B, top**). After refolding, a fully denatured band of 88 kDa became visible, more consistent with its theoretical size (**Fig. 5B, middle**). We also tested an EGFP fusion protein, human oxysterol-binding protein N-terminally fused with FLAG tag and EGFP, to find this fusion protein gave barely detectable fluorescence after the in-gel refolding despite the successful refolding of EGFP itself (**Supplementary Fig. S1**). This implies that the refolding efficiency of EGFP was diminished by the presence of the fusion partner protein, possibly through unproductive interaction between OSBP and EGFP polypeptides during in-gel refolding. Thus, the in-gel refolding would apply to GFP fusion proteins, but the GFP moiety requires higher folding efficiency than EGFP.

## 4. Conclusion

In this study, we developed a method to refold fully denatured GFPs in SDS-PAGE gels, which enables in-gel detection of fully denatured turboGFP-fused proteins. Shaking in Tris-Gly buffer with 1% αCD and 20% MeOH effectively removed SDS on the protein within 30 min, regenerating GFP fluorescence within the gel. The gel after the in-gel renaturation is compatible with total protein staining or immunoblotting without any additional steps. Thus, this method would offer a fast and easy alternative to the immunodetection of GFP-tagged proteins and should serve as a complementary method to the non-boiling or partial-denaturation SDS-PAGE [10]. It should be noted that the refolding efficiency can be affected by the presence of the partner proteins fused to the GFPs and also dependent on the GFP variant used. Although we showed that this protocol enabled in-gel fluorescence detection of fully denatured turboGFP-fused proteins, future studies are needed to clarify the exact scope of GFP variants and other fluorescent proteins amenable for in-gel refolding.

## Author contributions: CRediT

Misa Shiratori: Investigation, Formal analysis, Data curation, Writing -review & editing.

Rio Tsuyuki: Investigation, Resources, Writing -review & editing.

Miwako Asanuma: Investigation, Writing -review & editing.

Saki Kawabata: Investigation, Resources, Writing -review & editing.

Hiromasa Yoshioka: Investigation, Writing -review & editing.

Kenji Ohgane: Conceptualization, Investigation, Formal analysis, Data curation, Funding acquisition, Resources, Supervision, Project administration, Writing -original draft, Writing -review & editing.

## Acknowledgments

The authors wish to acknowledge the Institute for Human Life Science, Ochanomizu University, for the use of an iBright FL1500 imaging system (Thermo Fisher Scientific).

