## Supplementary Figure and Methods for "In-gel refolding allows fluorescence detection of fully denatured GFPs after SDS-PAGE"

(short title: Optimized in-gel refolding protocol for GFPs)

Misa Shiratori [a], Rio Tsuyuki [a], Miwako Asanuma [a], Saki Kawabata [a], Hiromasa Yoshioka [b], Kenji Ohgane [a,c,*]

[a] Department of Chemistry, Faculty of Science, Ochanomizu University, 2-1-1 Otsuka, Bunkyo-ku, Tokyo 112-8610, Japan

[b] Chemical Biology Research Group, RIKEN Center for Sustainable Resource Science, Wako, Saitama 351-0198, Japan

[c] Institute for Human Life Science, Ochanomizu University, 2-1-1 Otsuka, Bunkyo-ku, Tokyo 112-8610, Japan

### Table of contents

**Supplementary Figure**

**Figure S1:** Application of the refolding procedure to an EGFP-tagged OSBP stably expressed in HEK293 cells.

**Supplementary Methods**

**Megaprimer sequence 1:** GFPuv-to-sfGFP

**Megaprimer sequence 2:** OSBP-insertion

### Supplementary Figure

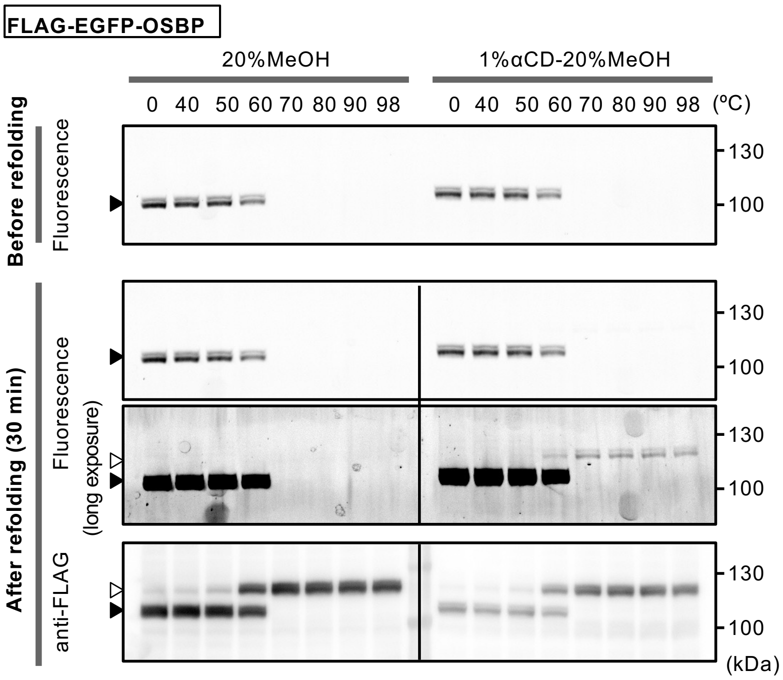


**Figure S1.** Application of the refolding procedure to an EGFP-tagged OSBP stably expressed in HEK293 cells. Aliquots of cell lysate mixed with Laemmli sample buffer were heated at the indicated temperature for 5 min and resolved on an SDS-PAGE with a 7.5% gel. The GFP fluorescence was imaged before and after the in-gel refolding with the Tris-Gly buffer supplemented with 1% αCD and 20% MeOH (30 min). A GFP image acquired with a longer exposure is also shown for the gel after refolding. Proteins on the gel were then transferred to a PVDF membrane, and the western blotting with anti-FLAG antibody was performed. The white and black arrowheads represent 120 kDa and 109 kDa, respectively.

### Supplementary Methods

**Megaprimer sequence 1.** GFPuv-to-sfGFP

The megaprimer below was used to convert GFPuv in the pGLO-high plasmid into superfolder GFP. The sequence covers a part of the GFPuv coding sequence and was used to introduce multiple mutations simultaneously by MEGAWHOP. The underlines denote mutations (S30R, Y39N, F64L, S65T, N105T, Y145F, I171V, and A206V) introduced to the GFPuv sequence.

CTGTC<u>AGA</u>GGAGAGGGTGAAGGTGATGCTACA<u>AAC</u>GGAAAGCTTACCCTTAAATTTATTTGCACTACTGGA
AAACTACCTGTTCCATGGCCAACACTTGTCACTACT<u>CTCACC</u>TATGGTGTTCAATGCTTTTCCCGTTATCC
GGATCATATGAAACGGCATGACTTTTTCAAGAGTGCCATGCCCGAAGGTTATGTACAGGAACGCACTATAT
CTTTCAAAGATGACGGG<u>ACC</u>TACAAGACGCGTGCTGAAGTCAAGTTTGAAGGTGATACCCTTGTTAATCGT
ATCGAGTTAAAAGGTATTGATTTTAAAGAAGATGGAAACATTCTCGGACACAAACTCGAGTACAAC<u>TTT</u>AA
CTCACACAATGTATACATCACGGCAGACAAACAAAAGAATGGAATCAAAGCTAACTTCAAAATTCGCCACA
AC<u>GTT</u>GAAGATGGATCCGTTCAACTAGCAGACCATTATCAACAAAATACTCCAATTGGCGATGGCCCTGTC
CTTTTACCAGACAACCATTACCTGTCGACACAATCT<u>GTC</u>CTTTCGAAAG

**Megaprimer sequence 2.** OSBP-insertion

The bold regions indicate the sequences that overlap with the template plasmid pReceiver-EGFP, and the underline indicates a linker sequence (TRTRPLE) inserted between EGFP and hOSBP.

**CACTCTCGGCATGGACGAGCTGTACAAG**A*CGCGTACGCGGCCGCTCGAG*GCTGCTACAGA
ACTTAGGGGAGTTGTCGGACCTGGACCTGCCGCCATTGCTGCTCTTGGCGGAGGCGGAGC
TGGACCTCCTGTTGTTGGAGGTGGCGGTGGAAGGGGAGATGCTGGTCCTGGATCTGGTGC
CGCTTCTGGAACAGTGGTTGCTGCTGCTGCAGGCGGTCCTGGACCAGGTGCTGGCGGAGT
TGCAGCTGCTGGACCAGCTCCTGCTCCACCAACAGGTGGAAGCGGAGGATCTGGCGCTGG
TGGATCTGGATCTGCCAGAGAAGGCTGGCTGTTCAAGTGGACCAACTACATCAAGGGCTA
CCAGCGGCGTTGGTTCGTGCTGTCTAATGGCCTGCTGAGCTACTACCGGTCCAAGGCCGA
GATGCGGCACACCTGTAGAGGCACCATCAATCTGGCCACCGCCAACATCACCGTGGAAGA
TAGCTGCAATTTCATCATCAGCAACGGCGGAGCCCAGACCTACCACCTGAAAGCCAGCAG
CGAGGTGGAACGGCAGAGATGGGTTACAGCCCTGGAACTGGCCAAGGCCAAAGCCGTGAA
AATGCTGGCCGAGTCCGATGAGTCTGGCGACGAGGAATCTGTGTCCCAGACCGACAAGAC
CGAGCTGCAGAACACCCTGAGAACCCTGAGCAGCAAGGTCGAGGACCTGAGCACCTGTAA
CGACCTGATCGCCAAGCACGGAACAGCCCTGCAGAGAAGCCTGAGCGAGCTGGAATCTCT
GAAGCTGCCCGCCGAGAGCAACGAGAAGATCAAGCAAGTGAACGAGCGGGCCACACTGTT
CCGGATCACCAGCAACGCCATGATCAACGCCTGCCGGGACTTCCTGATGCTGGCCCAGAC
ACACAGCAAGAAGTGGCAGAAGTCCCTGCAGTACGAGCGGGACCAGCGGATCAGACTGGA
AGAGACACTGGAACAGCTGGCCAAGCAGCACAATCACCTGGAAAGAGCCTTCAGAGGCGC
CACCGTGCTGCCTGCTAATACCCCTGGCAATGTCGGCAGCGGCAAGGATCAGTGTTGTTC
CGGCAAGGGCGACATGTCCGACGAGGACGACGAGAACGAGTTCTTCGACGCCCCTGAGAT
CATCACCATGCCTGAGAATCTGGGCCACAAGCGGACCGGCAGCAATATCTCTGGCGCCTC
CAGCGACATCAGCCTGGATGAGCAGTACAAGCACCAGCTGGAAGAAACGAAGAAAGAGAA
GCGGACTCGGATCCCCTACAAGCCCAACTACAGCCTGAACCTGTGGTCCATCATGAAGAA
CTGCATCGGCAAAGAGCTGAGCAAGATCCCCATGCCTGTGAACTTCAACGAGCCCCTGAG
CATGCTGCAGAGACTGACAGAGGACCTGGAATACCACGAACTGCTGGACAGAGCCGCCAA
GTGCGAGAACTCTCTCGAGCAGCTGTGTTACGTGGCCGCCTTTACCGTGTCCAGCTACAG
CACCACCGTGTTCAGAACCAGCAAGCCCTTCAATCCCCTGCTGGGCGAGACATTCGAGCT
GGACAGGCTCGAGGAAAACGGCTACAGATCCCTGTGCGAGCAAGTGTCCCACCATCCTCC
AGCCGCTGCTCATCACGCCGAGTCTAAGAATGGCTGGACCCTGAGACAAGAGATCAAGAT
CACCTCCAAGTTCCGGGGCAAGTACCTGAGCATCATGCCCCTGGGCACCATCCACTGCAT
CTTTCACGCCACCGGCCACCACTACACATGGAAGAAAGTGACCACCACAGTGCACAATAT
CATCGTGGGCAAGCTGTGGATCGACCAGAGCGGCGAGATCGACATCGTGAACCACAAGAC
CGGCGACAAGTGCAACCTGAAGTTCGTGCCCTACAGCTACTTCAGCCGCGACGTGGCCAG
AAAAGTGACCGGCGAAGTGACAGACCCCTCTGGCAAGGTGCACTTTGCTCTGCTCGGCAC
CTGGGATGAGAAGATGGAATGCTTCAAGGTGCAGCCCGTGATCGGCGAAAATGGCGGAGA
TGCCAGACAGAGAGGACACGAGGCCGAGGAATCCAGAGTGATGCTGTGGAAGAGAAACCC

TCTGCCTAAGAACGCCGAGAACATGTACTACTTCTCCGAGCTGGCCCTGACTCTGAACGC

CTGGGAAAGTGGCACAGCCCCTACCGATTCTAGACTGAGGCCCGACCAGCGGCTGATGGA

AAATGGCAGATGGGACGAAGCCAATGCCGAGAAGCAGAGACTCGAAGAAAAGCAGAGGCT

GAGCCGGAAGAAGAGGGAAGCCGAAGCCATGAAGGCCACCGAGGATGGCACACCCTACGA

TCCCTATAAGGCCCTTTGGTTCGAGCGGAAGAAAGACCCCGTGACCAAAGAACTGACCCA

CATCTACCGGGGCGAGTACTGGGAGTGCAAAGAGAAACAGGATTGGAGCAGCTGCCCCGA

CATCTTC**TAGCTCGAGTGCGGCCGCAACCCAGCTTTC**
